# Buffering of developmental noise provides a mechanism for heterosis in both polyploid and diploid hybrids

**DOI:** 10.64898/2026.05.27.728238

**Authors:** Gavin C. Conant

## Abstract

Using an abstract computational model of multicellular development, I show that the deleterious effects of gene expression noise on development are heavily buffered by increased ploidy, both from haploid to diploid cells and from diploid to tetraploid ones. Because the development of large multicellular organisms requires at least millions of individual cell divisions and many specifications of cell fate, it is likely impossible for large organisms to sufficiently control expression noise so as to prevent all noise-related errors in fate determination. However, for any given level of noise tolerance, cells of higher ploidy are less likely to suffer fate determination failure, potentially giving an explanation for the preference for larger organisms to have diploid somatic phases. Ploidy, however, is not the only potential mechanism by which differences in noise tolerance might influence hybrid vigor. Diploid hybrids can also display developmental robustness due to the removal of allelic correlations in expression noise. If we flip this perspective, noise-related disruption of development provides a neutral source of inbreeding depression, whereby sequence similarity throughout the genome induces correlated gene expression noise, reducing developmental robustness and pushing diploids back in the direction of haploid developmental fidelity.

## Introduction

Heterosis, or hybrid vigor, is the observation that crossing genetically distinct lineages can yield offspring with traits that are more desirable than those of either parental lineage [1, 2]. Discussions of it go back at least to Darwin, who devoted a book to the subject [3]. Those discussions tended to focus more on plants than on animals, likely because hybridization was more readily observed there [4, 5]. The early explanations for heterosis focused on the reduced frequency of double recessive genotypes in hybrids [6, 7]. In this view, because each population possesses a different set of mildly deleterious recessive alleles, crossing them will produce hybrid offspring with at least one fully-functional copy of each gene and a resulting increase in fitness. However, this explanation is likely incomplete for a few reasons. Different types of traits differ in their heterotic behavior [8], and the deficiencies seen in inbreed populations also vary by population [9]. Moreover, this masking of deleterious recessive alleles should be less evident in polyploid organisms, meaning they should be less subject to heterosis, a prediction that is not generally born out [7, 10, 11].

More recently, a more refined version of this hypothesis has focused on nearly neutral variation in gene expression. Such variation is due to small effects from many loci and has the effect of driving populations slightly off of fitness peaks in their gene expression. It is then argued that hybrids possess roughly the average of the expression of the parents. If each parent’s deviation from the optimal expression level were random, the average observed in the hybrids will generally be closer to that optimal level [12]. An attractive feature of this idea is the fact that it allows heterosis to derive from small effects at a large number of loci. Extending on this model, Okubo and Kaneko [13] developed a gene-regulatory network model of gene expression and evolved a number of populations to a gene expression fitness peak under the model. They then created fully homozygous and hybrid offspring from these models and assessed their fitness. The hybrid models showed both higher expression fitness and lower gene expression noise than did the homozygous models.

Importantly, hybridization is sometimes coupled to polyploidy. The resulting *allopolyploids* are potentially more fruitful than simple diploid hybrids for a few reasons: they do not require equal chromosome numbers, reproductive isolation may be immediate, and greater divergence between the hybridizing lineages may be tolerated [5, 14, 15]. The advantages of polyploidy-coupled hybridization are illustrated by the fact that allopolyploid plants were unusually likely to have been targets of domestication [16].

One feature of allopolyploidy and its contribution to heterosis that has been underappreciated is its effects on gene expression noise levels. Because gene expression is subject to stochastic effects due to the small numbers of molecules involved, the levels of molecules like mRNAs will vary even between genetically identical cells [17, 18]. Experiments in both yeasts and animals have shown that this expression noise decreases more or less directly with ploidy [19, 20]. This behavior can be understood as resulting from the close scaling of cellular volume and genome size [21]. This scaling will mean that large polyploid cells have similar cellular protein concentrations to diploids, but reduced noise due to their increased “sample size” of gene copies [22]. Unsurprisingly, stochastic models of gene expression show that transient drops in expression are more likely to be harmful to haploid cells than to diploid ones [23].

If heterosis needs to be considered in the context of expression noise and cellular ploidy, another factor that should also be included is development. The development of a multicellular plant or animal from a single cell is one of the most complex processes in biology, depending on a great range of processes including developmental timing, cell-to-cell communications, intracellular signaling and physical forces [24–26]. Many computational models of this process have been created, focusing on features such as cell-type differentiation and stem cell maintenance [27, 28], the morphological development of the physical embryo [29] and on integrating the genetic, cellular and morphological scales [30]. Curiously, these models have not yet been applied to hybrid vigor, despite the contribution of developmental traits like organism size to heterosis [6]. Hence, an analysis of the interplay of development, expression noise and ploidy is overdue.

When considering such models, the scale of the numbers involved becomes important. A typical human has on the order of 10 trillion cells [31] and hence must have undergone at least a similar number of cell divisions. Those cells are divided into at least 1,000 distinct types [32]. It is implausible to think that across this large number of divisions, expression noise could be so strongly controlled that it never contributes to differentiation failures, since that precision would imply fantastically small per-cell errors rates.

One place to look for evidence of a linkage between expression noise and developmental success might be in the distribution of life-cycle ploidies across organisms. Many organisms exhibit both haploid and diploid phases in their lifecycles, and the forces acting to determine the role of these phases can be complex [33]. However, there is a positive association between increasing size of the vegetative state and a tendency for that state to be diploid [33], just as one might predict if noise buffering improved developmental fidelity for larger organisms.

In this paper, I explore the effects of cell-type determination failures due to expression noise on a very simple computational model of organismal development. Increased ploidy can buffer developmental failures. Interestingly, increased diploid genetic diversity can also provide such a buffer if there are allelic noise correlations. Together, these results suggest that hybrid vigor and inbreeding depression might partly be shaped by the deleterious effects of intracellular noise on development.

## Results

### A simplified model of organismal development subject to cellular noise

I created an abstract model of organismal development that follows cell divisions on a balanced binary tree (Figure 1B). In it, the organism undergoes *t* cell type transitions to produce *2^t^* cell types. Because the organism also undergoes other cell divisions without type transitions, the total number of cells is *2^b(t+1)-1^*, where *b* is the number of cell divisions between each cell-type transition (*Methods*). I define the resulting organismal cell-type plan as the “programmed developmental outcome” or PDO.

**Figure 1.**
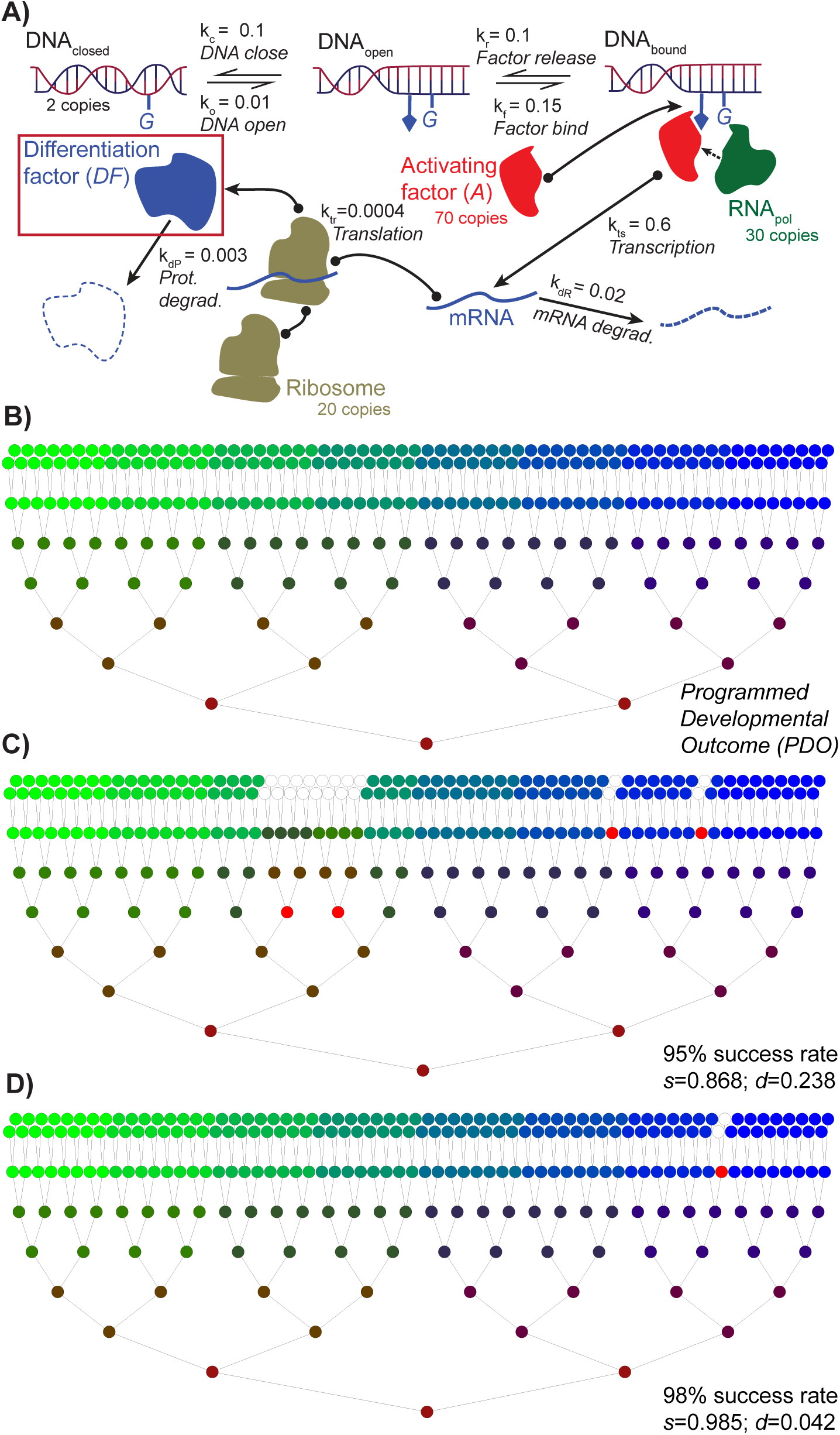
Modeling organismal development in the presence of cellular noise. **A)** A noisy model of gene expression. An external signaling factor *A* stochastically binds to a gene *G* in an open chromatin state and allows transcription. The resulting mRNA then undergoes competition between degradation and translation with a ribosome to produce a differentiation factor *DF*, which also has an intrinsic half-life. The DNA of *G* transitions back and forth between an open and closed state, giving rise to “burst” dynamics in mRNA production (*Methods*). Model parameters for the main model used are shown: I also tested randomized models with the same structure but differing parameters (*Methods*). **B)** A binary tree model of cellular differentiation. Starting with a single undifferentiated cell, cell divisions occur, and, at every *b* generations, the cells transition to a new cell type, up to *t* transitions. In the final state, there are *2^t^* cell types distributed across *2^b(t+1)-1^* total cells. This pattern defines the programmed developmental outcome (PDO). **C)** Cellular noise causes a failure of 1-R cell type transitions (red cells), resulting a simulated organism with fewer total cells due to apoptosis of the failed cells (white) and where the simulated organism does not match the PDO in the distribution of cell types. **D)** Increasing the developmental robustness to cellular noise reduces the difference between the simulated organism and the PDO.

To simulate the effects of cellular noise on this PDO, I embed a model of noisy gene expression inside of each cell undergoing a cell-type transition (Figure 1A). In this model, an external signaling factor *A* enters the cell at a fixed level. It binds upstream of a gene *G* only when that gene has an open chromatin configuration (*DNA_open_*). That binding can then induce the transcription of *G*. The resulting mRNA can then be degraded or translated into a “differentiation factor” (*DF*). The cell-type transition is deemed to be successful if the cellular particle count of *DF* is above a set threshold (see below). Noise in this system is due to several effects, including random mRNA and protein degradation, as well as the binding of *A* and the RNA polymerase to *G* and of the ribosome to the mRNA. However, the primary driver of noise in *DF* is the transition between open and closed DNA, which occurs on a timescale of seconds [34–38].

We can explore the role of such expression noise on the success of development by defining a per-cell transition success rate *R*. If *R=0.95*, then there is a 95% chance that any particular cell transition will succeed and a 5% chance of failure (red cells in Figure 1B&C). The cellular noise model was implemented with the stochastic simulation algorithm of Gillespie [39], as implemented in the Stochkit2 package [40] (*Methods)*. From these simulations, the distribution of *DF* counts is obtained and the cutoff value for *DF* computed from the value of *R*. Because development is occurring on a binary tree, this definition has two implications. First, the larger *R,* the fewer failures in the PDO. Second and more important, failures will on average occur late in development, because there are many more cell transitions late in the process (Figure 1B&C). The model assumes that any cell that fails a transition will pass an invalid cell state to its offspring—I then impose an apoptosis step on the final organism, such that any cells that are not in the correct developmental stage are killed (white cells in Figure 1B&C). I define two metrics on the success of development for a particular simulated organism: *d*: the distance between the set of cell states in the PDO and the simulation and *s:* the size (in number of cells) of the simulated organism relative to the PDO (*Methods*).

### Autopolyploidy yields model organisms with greater fidelity to the PDO

I used this modeling framework to explore the effects of a simulated autopolyploidy on developmental success. To do so, I simply increased the number of copies of *G* in the noise model from 2 to 4. To make the simulations comparable to the diploid version, I reduced the affinity of *A* for *G* such that the polyploid model produced the same *average* number of *DF* molecules (*Methods*). This approach mimics an autopolyploidy where the increase in DNA copy number is precisely balanced by a doubling of cellular volume [41], yielding unchanged cellular concentrations. I estimated the mean cutoff value for *DF* at a given value of *R* in the simulated diploids. In other words, if, when *R=0.95*, 95% of the simulations had *DF=*142 particles or more, I then assumed that transition would occur correctly if *DF≥*142 in the polyploid cells as well. However, because the polyploids are producing *DF* from 4 loci rather than 2, the intrinsic noise in *DF* is reduced. As can be seen in Figure 2, at any given success rate in the diploids, the polyploid model gives better average fidelity to the PDO.

**Figure 2.**
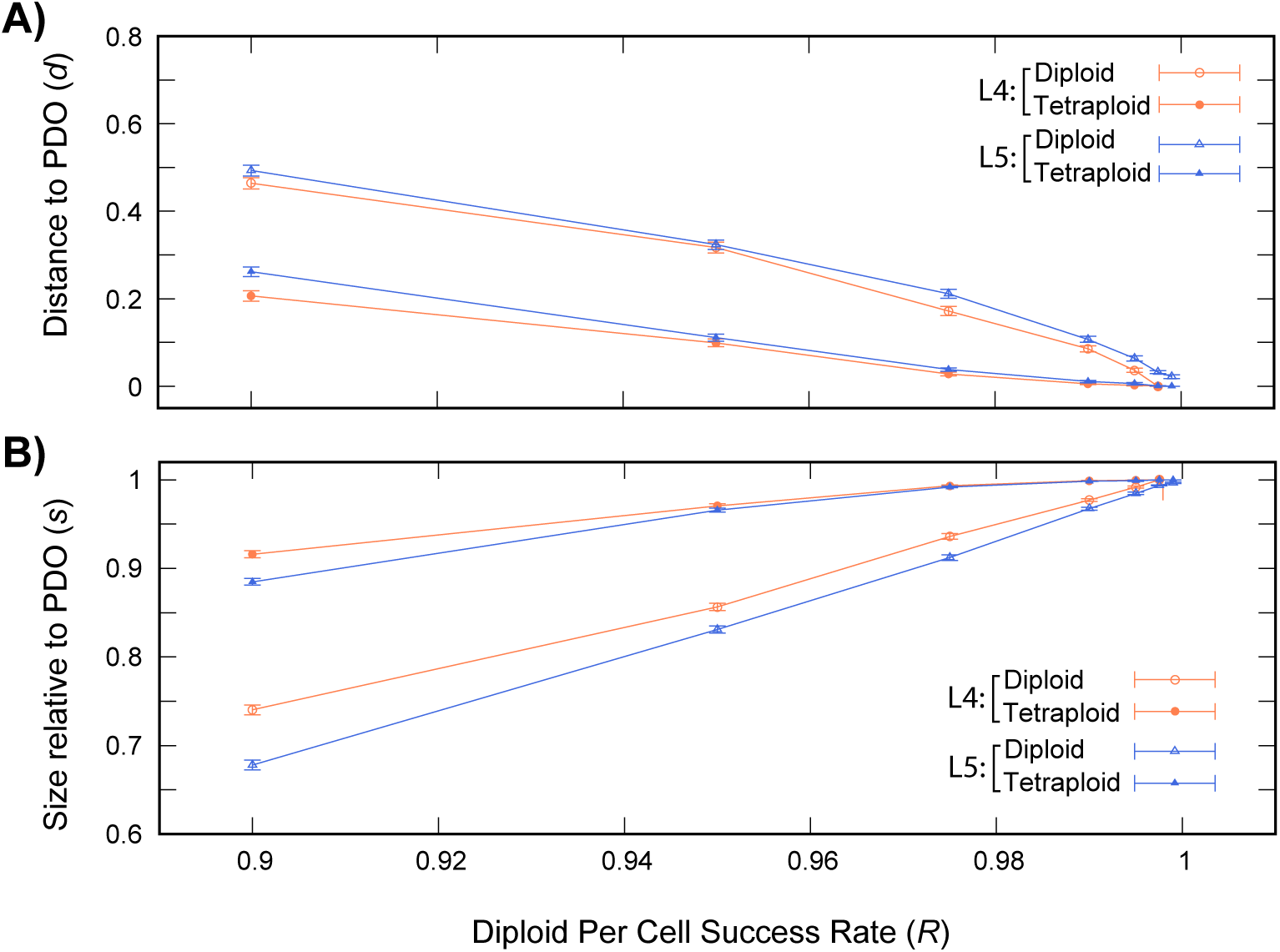
Polyploidy increases developmental fidelity. **A)** Comparing simulated organisms with 4 (L4) and 5 (L5) total cell-type transitions, corresponding to 512 and 2048 final cells, respectively. On *x* is the per-cell transition success rate *R.* As *R* approaches 1.0, the number of transition failures in the cells of the model falls. On *y* is the Euclidian distance *d* between the average simulated developmental outcome (over *n=*200 simulations) compared to the PDO. Cases where *d=*0 correspond to a distribution of cell types identical to the PDO. Open points show the diploid outcomes, closed points the polyploid ones. The polyploid models show higher fidelity across most per-cell success rates. For the L4 models, the analysis was stopped at *R*=0.9975, since at that point the expected number of transition failures is less than one over the number of transitions in the diploid model. Error bars give the standard error of the sample mean [63]; namely the sample standard deviation over 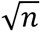. **B)** Diploid and polyploid developmental simulations show different fidelity to the PDO. On the *y*-axis is the ratio of the size of simulated organism compared to the PDO (average of 200 simulations). When *s=1*, the simulated organism has the same number of cells as the PDO (i.e., no apoptosis occurred in development). Error bars give the standard error of the sample mean [63]. The polyploid models again show greater fidelity. For L4, all comparisons except *d* and *s* at *R*=0.9975 (where the diploid matched the PDO) show significant differences between the diploid and polyploid (*P*<10^-5^; unpaired *t*-test*. Methods*). All comparisons between the diploid and polyploid for L5 showed significant differences in mean *s* and mean *d* (*P*<10^-5^; unpaired *t*-test*. Methods*).

### The developmental buffering effects of polyploidy are seen across a range of model types

To be sure that the results in Figure 2 were not an artifact of that particular model, I explored the developmental behavior of polyploids under a range of assumptions. I defined a robust developmental model where cells can recover from a transition failure in the next round of cell divisions, such that such failures do not invariable force apoptosis. I also treated polyploidy as resulting in a doubling of the production rate of *DF* (rather than the more concentration-like approach above). To do so, I doubled the cutoff of *DF* seen in the diploid when constructing the corresponding polyploid (*Methods*). Finally, I constructed 100 random noise models with the structure of Figure 1A but differing kinetics and estimated the intrinsic noise in *DF* across these models (*Methods*). I then selected the model with the lowest noise, that with the highest noise and that with the midpoint level of noise. I compared the effects of polyploidy on each of these three random models, finding that polyploidy resulted in greater developmental fidelity in all three cases (Figure 3).

**Figure 3.**
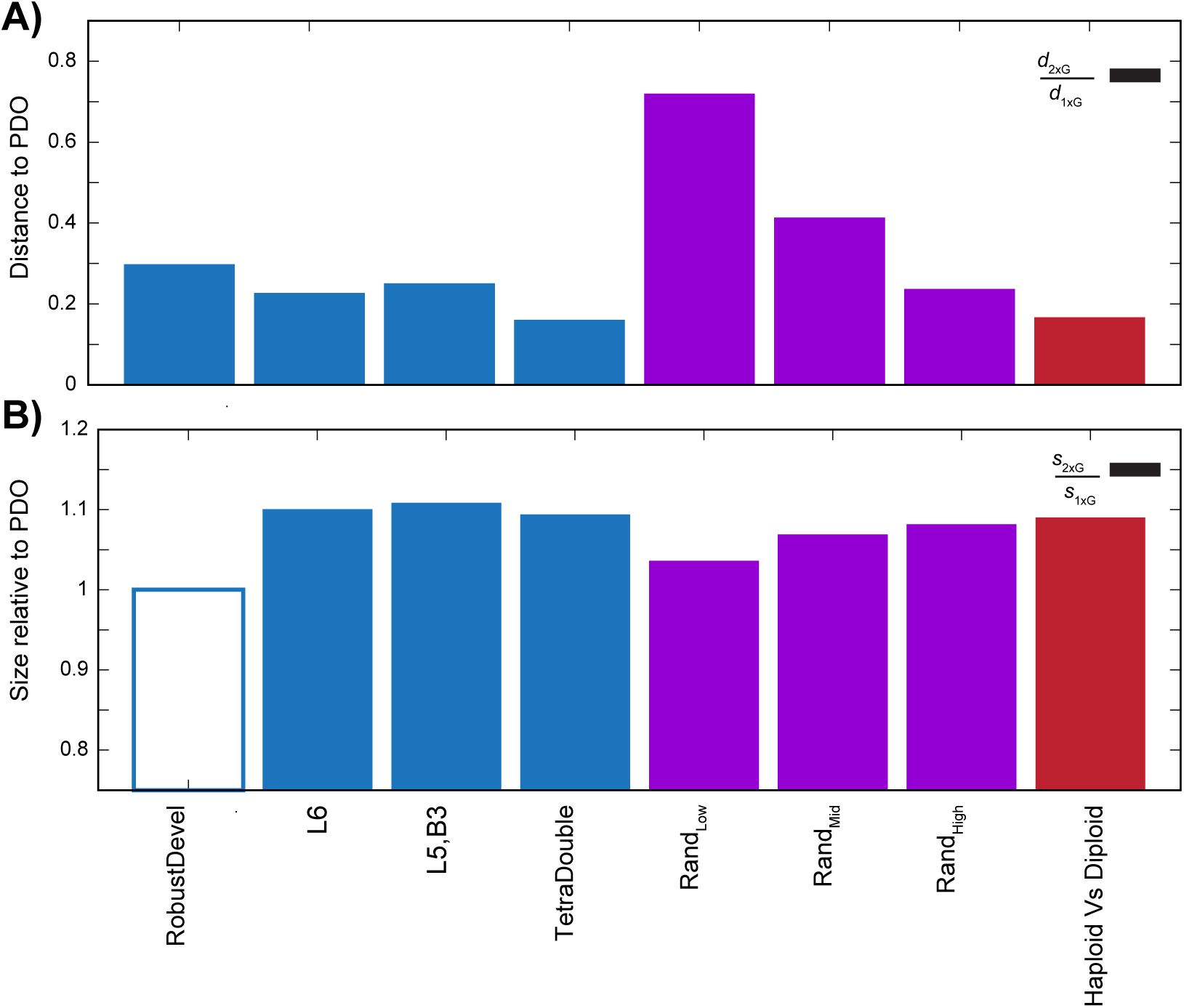
The increased developmental fidelity of polyploids is consistent across a range of models (*Methods*). I compared a model robust to transition failures (“RobustDevel”), a larger version of the model in Figure 2 with 8192 final cells (“L6”), a model with more intermediate cell divisions (*b=3* rather *b=2* in Figure 2, meaning a total of 131,072 final cells) between transitions (“L6,B3), and model where polyploidy induces a doubling in cellular *DF* levels (“TetraDouble”). I also compared three random models with increasing levels of noise (“Rand_Low_,” “Rand_Mid_” and “Rand_High_”). Finally, I compared a diploid model to a haploid one. **A)** Observed effects of polyploidy at *R*=0.975 on the distance *d* to the PDO. Shown in each bar is the ratio of the polyploid (or diploid) value of *d* (*d_2xG_*) over that of diploid/haploid (*d_1xG_*). Values less than one correspond to greater fidelity to the PDO in the polyploids. All differences in mean *d* between the diploids and polyploids are significant (*P*<10^-5^, unpaired *t*-test; Methods) **B**. Effects of polyploidy on final organism size. Shown in each bar is the ratio of the polyploid (or diploid) value of *s* (*s_2xG_*) over that of diploid/haploid (*s_1xG_*). Values greater than one correspond to greater fidelity to the PDO in the polyploids. No significant difference between the diploid and polyploid developmental models were seen in *s* for the robust model (“RobustDevel;” box outlined; *P>*0.5; unpaired *t*-test; *Methods*) because this model does not induce apoptosis unless there are sequential failures in transition. Instead, failures result in an offspring with a differing distribution of final cell types (*d* above). All other comparisons showed significant differences in mean *s* between the diploids and polyploids (*P*<10^-5^, unpaired *t*-test; Methods).

### Diploid cells also have developmental advantages over haploids

The arguments above should apply equally to haploid verses diploid cell development, and so I compared a model with one copy of *G* to one with two copies using the same approach. Once again, increasing the gene copy number reduces cellular noise and stabilizes development (Figure 3).

### Developmental resilience and non-polyploid hybrids

Since diploid hybrids also experience heterosis [3, 6, 7], an obvious question is how developmental robustness is effected by hybridization between lineages with different noise behavior. As a baseline, I used the random models just mentioned and selected 20 pairs of those models such that the two members of a pair had similar variances in their *DF* levels. I then hybridized these pairs. Unsurprisingly, the estimated cutoffs in *DF* (at *R*=0.975) for the hybrids was intermediate between their parents (Supplemental Figure 1). In two cases, however, the developmental outcome of the hybrid was significantly improved over one parent and in one case it was improved relative to both (Supplemental Figure 1). This result is consistent with our prior work that emphasizes the nonadditive nature of gene expression inheritance [42], though it is difficult to generalize the mechanism involved with such a simplified expression model.

The simplicity of the expression models probably also makes observing any strong heterotic behavior unlikely. In particular, all of the noise models used have independent transitions between open and closed DNA for the alleles of *G* and independent binding of activation factors and polymerases to those alleles. Hence, the only coupling between alleles is from weak effects such as dilution of the level of *A* through its binding to one promotor. Unsurprisingly, when I computed the correlation between the level of *DF* produced by each allele of *G*, they were uncorrelated in both the individual parents and in the hybrids (−0.01<Pearson’s *r*<0.01).

In real organisms, patterns such as common DNA methylation sites may actually tend to produce correlations in the chromatin state of identical genes. To mimic this behavior in the model, I added a reaction that allows each allele to open or close the other allele’s chromatin state when the two states are different (*Methods*), forming model *M_corr_*. Doing so can induce a relatively high correlation between the level of *DF* produced by the two alleles (Figure 4).

**Figure 4.**
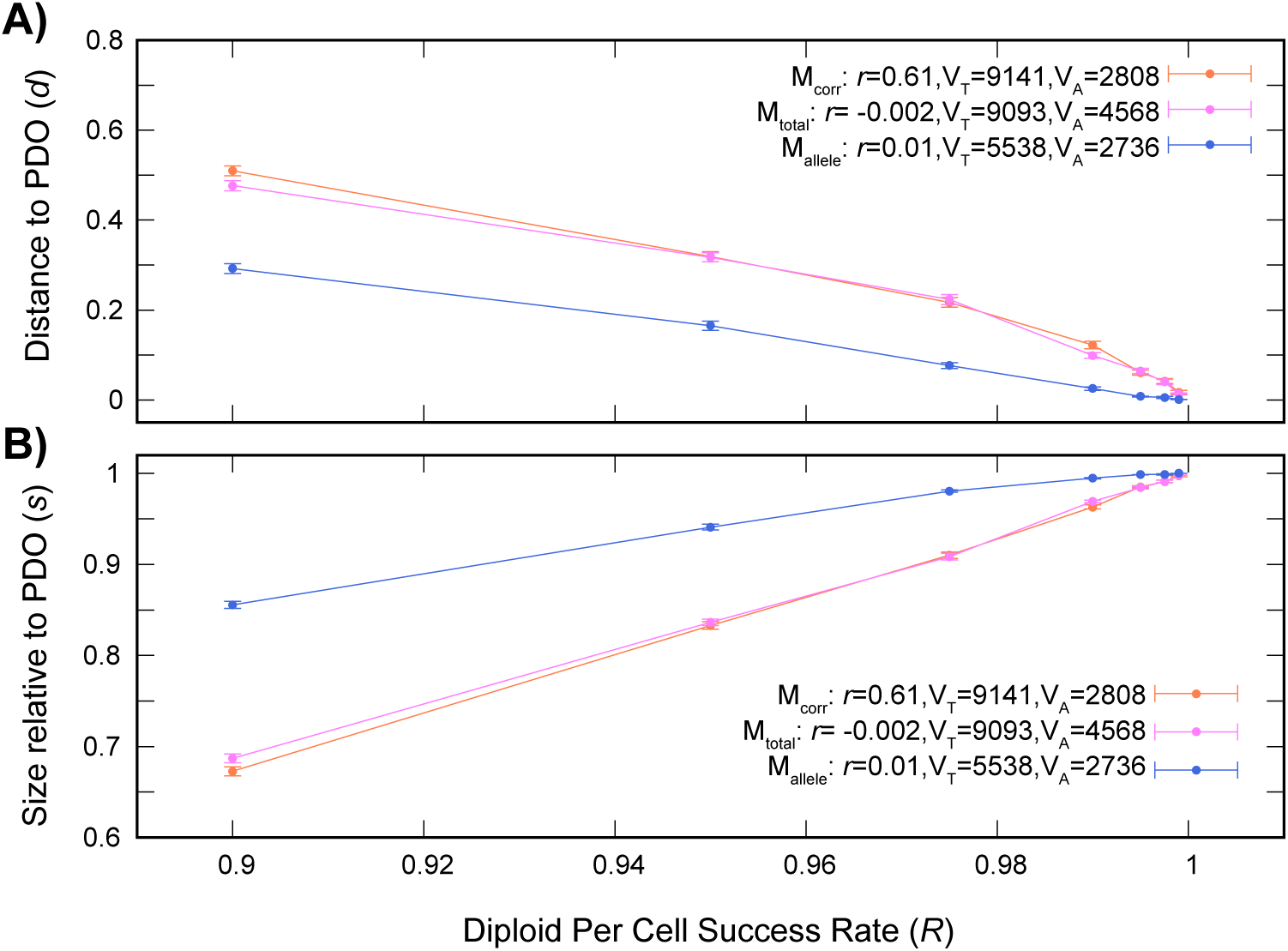
Allele expression correlations in diploids produces no apparent excess of developmental failures but hybrids lacking this correlation can show higher developmental fidelity. Comparing a model with a correlation in allele expression (*M_corr_*) to one with the same total variance in the level of *DF* but no correlation (*M_total_*) produces very similar error profiles. However, a model with the same level of allelic variance (*M_allele_*) that might occur through hybridization breaking the allelic correlations, gives reduced distance to the PDO. **A)** As transition success rate *R* increases, the distance *d* to the PDO decreases in all cases, but the mean *d* for *M_allele_* is always closer to the PDO than is *M_corr_* (*P<*10^-5^; unpaired *t*-test; *Methods*). Models *M_corr_* and *M_total_* have generally indistinguishable means, except for at *R*=0.9 and *R*=0.99 (*P=*0.03 in both cases; unpaired *t*-test). **B)** The size relative to the PDO is also similar for *M_corr_* and *M_total_* but different for *M_allele_*, with *M_allele_* is always closer to the PDO than is *M_corr_* (*P<*10^-5^; unpaired *t*-test): *M_corr_* and *M_total_* differ only at *R*=0.9 (*P=*0.02; unpaired *t*-test).

Because the developmental model responds only to the total *DF* level, I can create an uncorrelated model with the same total variance in *DF* as in *M_corr_* (model *M_total_*). This model produces a developmental error profile essentially indistinguishable from *M_corr_* (Figure 4). However, one can also measure the variability of each allele’s *DF* product individually. If the cell has evolved to minimize noise even in the presence of allelic correlations, that selective force should be evident in a low value of this allelic variance. I hence next created an uncorrelated model *M_allele_* where the allele-specific variance in *DF* was the same as that seen in *M_corr_* (*Methods*). Because the two alleles in *M_allele_* are independent and uncorrelated, the total variance in *DF* for this model is less than that total in *M_corr_*. As a result, when I compare the developmental stability of the three models, *M_allele_* has greater fidelity to the PDO than either of the other models (Figure 4). These effects are not specific to the model used but are also seen in when the allelic correlation is lower and in three random models of differing variability (Supplemental Figure 2).

## Discussion

There is a growing appreciation that any understanding of the evolution of gene expression must incorporate the existence of noise [13, 22, 43–45]. For instance, features of the eukaryotic transcriptional regulatory network such as the overabundance of feedforward loops [46] can be (partly) attributed to the need to minimize expression noise [47–49]. It is also apparent from both experiment and theory that noise scales inversely with ploidy [19, 20, 22, 23].

Existing work on the scaling of noise with ploidy has focused on effects inside cells [13, 23, 43]. A natural question that follows is whether expression noise has effects in multicellular organisms beyond those known from single cells. One of the more obvious places to look for such effects would be in development.

From the perspective of a single cell, noise can be viewed as random temporal deviations from the expected gene expression program, meaning that it will often have limited effects so long as average protein levels are appropriate [22]. Development, however, is a case where this simplified view is unlikely to be valid, as development is critically dependent on timing [50], meaning that noise-related reductions in expression could be particularly harmful there. In this vein, it is perhaps interesting that large multicellular organisms seem to prefer to have their somatic forms to be diploid [33], as one might predict if diploidy better buffered expression noise.

The results above extend on this hypothesis, albeit under a number of simplifying assumptions. The model of development used is obviously quite artificial, and the noise model includes only a single gene under very simple regulation. However, one should notice that the coupling of the cellular and developmental models is not strong: the results really depend only on there being some source of internal noise in the cells that drives differentiation failure at some low rate. The entities in Figure 1B-D also need not be thought of as precisely representing individual cells—they represent points in development where the fates of lineages diverge. Caution is also needed in interpreting the above results because the kinetic expression models used do not map naturally to genetic differences between individuals. The cell also has many noise buffering mechanisms not present in those models [48, 49]. Indeed, cellular noise is not always detrimental—there are examples where organisms use noise for robustness or adaptability, including in development [51–53].

Nonetheless, there are two key insights provided in this modeling framework. The first is that the value of higher ploidy in stabilizing processes such as development has been underappreciated. It is known than genes involved in developmental processes are over-retained in duplicate after plant polyploidies [54, 55]. I propose that part of the reason for this over-retention is that the noise-buffering effects of polyploidy do not require the retention of the entire genome in duplicate. Instead, the effects seen above could be achieved through the retention in duplicate of only the developmental factors involved. Some evidence for this idea is seen in the post-polyploidy evolution of the zebrafish genome. In that species, there is a deficit of retained duplicate genes from the teleost genome duplication expressed at the earliest developmental periods [56]. These early phases, where there are relatively few cell-type transitions, are characterized by the use of maternal mRNAs rather than those from the developing embryo [57]. But, the later phases, where the need to buffer noise should be higher, are instead characterized by an *excess* of retained duplicates from the polyploidy [56]. It is also worth considering whether the ability of polyploids to become established at times of environmental stresses [58] might owe something to the developmental buffering discussed here.

The second insight from the analyses above is that genetic diversity alone can also increase developmental robustness, at least under the slightly contrived modeling scenarios used. Even simple diploid hybrid models at least occasionally have improved outcomes over their parents, though I am reluctant to overstress this result, given the simple expression model used. More generally, to the degree that genetically similar alleles show correlations in their noise profiles, those diploid individuals will be less successful in buffering developmental errors because when one allele is abnormally expressed, it is more likely that the other one will be as well. Such correlations could arise, for instance, if allelic copies with similar sequences display similar interactions with chromatin modifying enzymes [59, 60].

An appealing feature of this second type of developmental robustness is that it offers a means by which the deleterious effects of inbreeding can arise in genomic regions evolving neutrally. In this view, some of the effects of inbreeding (and the benefits of hybridization) arise because increasing genomic similarity gives rise to increasing similar noise profiles between the two allelic subgenomes, even though individually those subgenomes have normal noise profiles. Bringing together differing genomes will reduce the correlated expression noise and give rise to individuals with greater developmental robustness. Whether this concept, which is quite reasonable from a modeling perspective, represents an important force in genome evolution in real organisms is worth testing.

## Methods

### A computational model of organism development

To assess the role of expression noise on development, I created a simplified model representing the development of an organism with a defined number of cells, subdivided into a specified number of cell types (Figure 1). The model defines a number of transitions *t* and a number of cell divisions between transitions *b*. Starting with a single cell at developmental level 0, the model moves through *b* generations of binary divisions then undergoes a transition until the total number of desired transitions is reached, after which *b* further divisions occur. Each cell has two cell-fate identifiers: its developmental level *l* (0≤*l*≤t) and its cell type *c* within that level (0≤*c*<*2^l^*). When a transition occurs, the cell produces two daughter cells at level *l+1*. The daughters’ value of *c* is determined by whether their parent cell was on the left or right side of the transition to state *c*, since the number of cell types doubles at each transition (Figure 1B).

Hence, the developmental profile involves *t+1* layers, each of size *b* generations. The final number of cell types in the model is therefore *2^t^* and the total number of cells in the ideal final organism is *2^b(t+1)-1^*, accounting for the *t+1* layers involved with the initial generation having had only a single cell in it. The number of cells of each type is thus

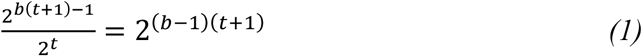

### Cellular noise model applied to the development model

This model produces the binary tree of Figure 1B, corresponding to the “programmed developmental outcome” (PDO). I now assume that, in the simulated organisms, this PDO is subject to noise. I use the simplified noise model outlined in Figure 1A. It assumes that there is some activating factor *A* that is present in the cell at a fixed concentration not subject to noise. This factor serves as a transcriptional activator for a gene *G*, allowing the RNA polymerase to be recruited to *G* and transcription to occur. However, binding, recruitment and transcription can only occur if *G* is in a transcriptionally accessible state. For mammalian cells, genes spend perhaps 90% of their time in an inaccessible state and transition between the two states on a timescale of tens of seconds [34, 35]. This transcriptional program tends to produce mRNA in “bursts” [36, 37]. Those mRNAs can then either be translated or degraded. The final protein product is a differentiation factor (*DF*) that drives the cell to correctly differentiate if it is present at sufficient levels. The mRNA half life is assumed to be on the order of minutes [61] and the protein half-life approximately 10 times longer [62]. This model yields a pattern of “noisy” mRNA levels with more stable levels of the *DF* [38]. The stochastic simulations from this model were run using the tau_leaping_exp_adapt_serial tool of the Stochkit2 package [40]. To avoid transient effects due to the models starting with no *DF*, I ran the development simulations for 3000 timesteps and sampled the DF state at that final timepoint.

Hence, our base model embeds a stochastic gene expression model inside a developmental model, with the level of *DF* at the transition point deciding if the cellular correctly transitions to two daughter cells with correct cell identities. When a transition fails (red cells in Figure 1B&C), the daughters inherit the level *l* of the parent, and when the final developmental stage is reached, any cells that have not reached *l*=*t+1* are assumed to undergo apoptosis (white cells in Figure 1B&C). To decide when a transition fails, I define *R*: the per-cell success rate of the developmental program. If *R=0.95*, then any cell whose level of *DF* is in the top 95% of the values seen in the stochastic simulations is deemed to have transitioned correctly, meaning that *R=0.95* corresponds to a 5% per-cell failure rate in transitions.

### Metrics on the success of development

I defined two measures of the deviation between the “perfect” PDO and a simulated developmental outcome under cellular noise. First, I defined on the outcome a vector *V* of length *2^t^* where each element is the number of cells of that type in the outcome. For the PDO, each element of *V^PDO^* is equal to *2^(b-1)(t+1)^* (in other words, the total number of cells divided by the number of cell types, equation 1). Because of failures in differentiation, the vector for a simulated outcome *V^S^* will have fewer cells of some types. I used the Euclidian distance *d* between *V^PDO^* and *V^S^* as a measure of the success of development:

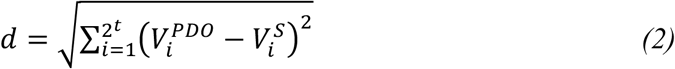

The second metric is simply the length or size of the final organism *s*, relative to that of the PDO, which is defined to have size *2^b(t+1)-1^*: any apoptosis in the simulated outcome will reduce its size relative to the PDO.

### Modeling ploidy changes in the simulated developmental outcomes

I now sought to model how a change in ploidy, either from haploid to diploid or from diploid to tetraploid, would affect the error profile of the developmental program. In the stochastic simulation framework of Stochkit, concentrations are not explicitly represented: Instead reactions occur based on particle numbers and relative reaction rates. In such a model, if I double the number of particles of *G* from 2 to 4 for a tetraploidy or from 1 to 2 for the transition from haploid to diploid while leaving all other factors in the model constant, that will double the production rate of *DF*. Since I want to assess the effects on developmental error rates, this change in *DF* levels is undesirable. I therefore reduced the binding rate constate *k_f_* of *A* for each of the copies of *G* (Figure 1A) such that the polyploid model produces the same average level of *DF* as the diploid model. I did so by repeatedly running the stochastic simulation with differing values of *k_f_* until the mean *DF* levels in the polyploid model converged to those of the diploid model. This procedure resulted in values of *k_f_* for the polyploid model of approximately ½ of that for the diploid model, as one would predict under the assumption of a 2-fold dilution of binding from a doubling of cellular volume. I then simulated the effects of polyploidy by using the value of *DF* corresponding to a particular value of *R* from the diploid model as the threshold for failure in the polyploid model as well. If the polyploid model has lower cellular noise levels due to the extra gene copies, that will be reflected in fewer cells failing to transition, which can be detected by comparing *d* and *s* to those seen from the diploid models (Figure 2). The mean values of *d* and *s* for the diploid and polyploid simulations were compared using two-sample *t-*tests [63], computed with R version 4.5.2 [64]. The same approach was used to compare the haploid and diploid models using the haploid model as the basis (Figure 3). For completeness, I also confirmed that allowing the *DF* concentrations to double in the polyploid model and doubling the threshold seen in the diploid model for the comparisons gave effectively the same results (see Figure 3).

### Noise model families

To be sure that these results were not idiosyncratic to the particular model parameters in Figure 1A, I created a family of 200 randomized versions of that model that differed in their 8 kinetic rates. The range of rate values used is shown in Supplemental Table 1. These random models were all optimized to produce an average of 200 molecules of *DF*: the model of Figure 1A produces 238.3 molecules of *DF* on average. For each model, I computed the variance of the *DF* and retained the 100 models with the lowest variances (i.e., the 100 least noisy models). To assess the role of intrinsic model noise on the effects of polyploidy, I simulated polyploidies for the least-variable model, the most variable model and the midpoint model (Figure 3). Supplemental Figure 1 shows values of *d* for 40 parental models used in hybridization experiments (see below): when *R* is constant, all of these models occupy a narrow range of *d* values.

### Model hybridization

A main value of the random models just described is that they allow the exploration of the effects of hybridization. I implemented hybridization by treating the eight rate constants in the model of Figure 1A as specific to each lineage, such that the hybrid inherits a copy of *G* from parent 1 (*G_1_)* with its inherent 8 rates, and a *G_2_* copy from the other parent with another set of constants. However, the activation factor (*A*) and the polymerase and ribosome are assumed to function interchangeably other than their different rate constants and the resulting *DF*s are also assumed to be interchangeable. I selected 20 pairs of models with similar variances in *DF* and created hybrid models. For each such pair, I simulated 800 developmental outcomes for the two parents and the hybrid using *R=*0.975.

### Models with correlation between allelic copies of the DF

I next made versions of the diploid models that allowed me to independently track the *DF* molecules made by the two alleles of *G* in the model. Unsurprisingly, the levels of the two versions of *DF* (from the two different alleles) were effectively uncorrelated after a transient “burn in” period of 20% of the simulation run time (*Results*). To create a model *M_corr_* with correlations between the two alleles, I added an interaction between the DNA state of the two genes, such that if one gene was in the open state and one in the closed, there was a rate at which the open gene could open the chromatin of the closed one and, at the same rate, the closed gene could close the chromatin of the open gene. This correlation in chromatin state creates correlations in the levels of *DF* created from those two alleles.

I next needed uncorrelated models to compare this model to. Firstly, I computed the total variance in *DF* seen in *M_corr_*. I then removed the interaction reaction to create an uncorrelated model. One can tune the variance in this model by scaling both *k_o_* and *k_c_* by a constant factor *c*. Increasing *c* reduces the transition time between open and closed DNA states, reducing the variance in *DF*; lowering *c* has the opposite effect. I thus tuned the uncorrelated model *M_total_* to have the same total variability in *DF* as seen in *M_corr_*. As expected, these two models show effectively identical behavior in the developmental simulations (Figure 4), since those simulations only consider the total *DF* levels. I then measured the *allele-specific* variance in *DF* in *M_corr_* and created a new model *M_allele_* where each allele of *G* produces *DF* with the same variability as the two alleles in *M_corr_* but with no correlation between those two alleles. This same analysis was applied to a model with a lower conversion rate between the open and closed state and for the three random models used above (Supplemental Figure 2).

## Acknowledgements

This work was supported by U.S. National Science Foundation grant NSF-DEB-2241312.

## Data availability

Expression models and scripts for their analysis are available from figshare (https://doi.org/10.6084/m9.figshare.32273766). The developmental simulation code is available from GitHub (https://github.com/gconant0/DevelopNoiseSim).

