## Supplementary material for "Buffering of developmental noise provides a mechanism for heterosis in both polyploid and diploid hybrids": Supplmental Tables and Figures

**Supplemental Table 1:** Noise model parameters in the base and random models

| **Rate**[a] | **Base model value**[b] | **Min. rand value**[c] | **Max. rand value**[c] |
|---|---|---|---|
| $k_{dP}$ | 0.003 | 0.0027 | 0.052 |
| $k_r$ | 0.1 | 0.025 | 0.58 |
| $k_o$ | 0.01 | 0.013 | 0.063 |
| $k_f$ | 0.15 | 0.032 | 1.04 |
| $k_{tr}$ | 0.0004 | $9.7 \times 10^{-5}$ | 0.0023 |
| $k_{dR}$ | 0.02 | 0.0067 | 0.11 |
| $k_{ts}$ | 0.60 | 0.13 | 3.32 |
| $k_c$ | 0.1 | 0.016 | 0.61 |

a: Model kinetic constants from the model scheme shown in Figure 1A. These values correspond to relative event rates in the simulations.

b: Constant value used in the model of Figure 1A.

c: Minimum and maximum value of that kinetic constant seen across the 100 optimized random models with lowest overall noise levels (*Methods*).

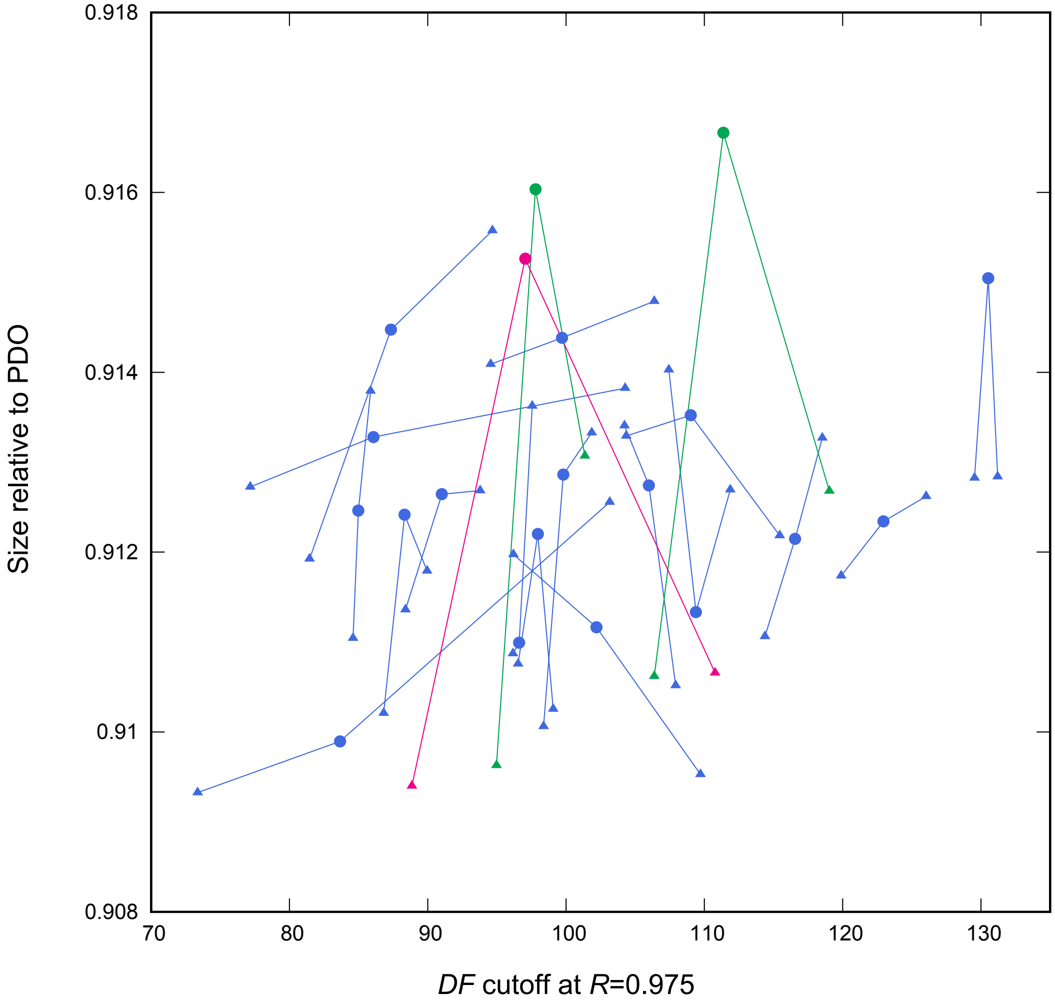


**Supplemental Figure 1**. Hybrid models (circles) compared with their parents (triangles connected by lines). On *x* is the cutoff value of *DF* that gives a success rate of *R*=0.975. In all cases, the *DF* cutoff for the hybrid is intermediate to that for the two parents and statistically different from each ($P<0.02$, unpaired *t*-test. *Methods*). On *y* is size relative to the PDO for all three models (*s,* 800 simulations, *Methods*). In two cases, the *s* of the hybrid was significantly higher than that for one parent (green; $P<0.02$, unpaired *t*-test. *Methods*) and in one case it was significantly higher than both parents (magenta; $P<0.02$, unpaired *t*-test. *Methods*).

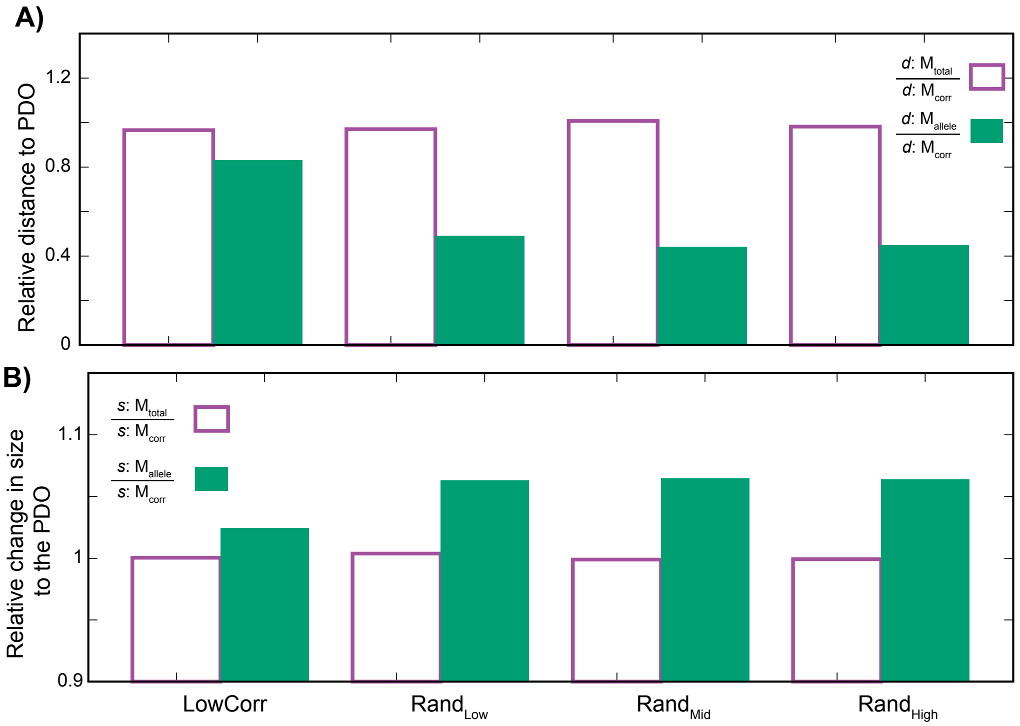


**Supplemental Figure 2**. Models with lower allelic correlations and random models with correlated allelic expression behave similarly to uncorrelated models with similar total variance in *DF* levels. However, models with similar allelic variance levels but no allelic correlations show enhanced fidelity to the PDO. For the original model and the three random models with low ("$\text{Rand}_{\text{low}}$") medium ("$\text{Rand}_{\text{Mid}}$") and high ("$\text{Rand}_{\text{High}}$") intrinsic variation in *DF*, I created a correlated version of each model ($M_{corr}$) where the chromatin state of the two alleles of *G* is correlated. For the "LowCorr" model, this approach yielded a model with a Pearson's correlation between $DF_1$ and $DF_2$ of $r$=0.15, compared to $r$=0.61 in Figure 4. I created a second model ($M_{total}$) where the total variance in *DF* was the same as in $M_{corr}$ but without allelic correlations. Finally, I created a third model ($M_{allele}$) where the individual variances in the two alleles was the same as in $M_{corr}$ but without any correlation in the state of the alleles of *G*. I then made 200 simulations of the developmental process ($t$=5; $b$=2) of the four models. I compared the four models for their similarity to the PDO for both *d* and *s*. **A)** Comparing the distance to the PDO between $M_{total}$ and $M_{corr}$ (left) and $M_{allele}$ and $M_{corr}$ (right) across the low correlation and three random base models (*x*-axis). In all four cases, the value of *d* is indistinguishable between $M_{total}$ and $M_{corr}$ (outlines, $P>0.05$; unpaired *t*-test. *Methods*). The $M_{allele}$ model is always closer to the PDO than is $M_{corr}$ ($P<0.01$; unpaired *t*-test. *Methods*). **B)** Comparing the three models in size relative to the PDO. In all four cases, the value of *s* is indistinguishable between $M_{total}$ and $M_{corr}$ (outlines, $P>0.05$; unpaired *t*-test. *Methods*). The $M_{allele}$ model is always closer in size to the PDO than is $M_{corr}$ ($P<10^{-5}$; unpaired *t*-test. *Methods*).
